# Laboratory-acquired mutations fall outside the wild-type alleleome of *Escherichia coli*

**DOI:** 10.1101/2022.04.04.487056

**Authors:** Edward Catoiu, Patrick Phaneuf, Jonathan Monk, Bernhard O Palsson

**Author notes:** **Corresponding Author** Correspondence to Bernhard O Palsson.

## Abstract

Inexpensive DNA sequencing has led to a rapidly increasing number of whole genome sequences in the public domain. Natural sequence variation can now be assessed across a large number of sequenced strains of a bacterial species, resulting in the definition of the wild-type *alleleome* (the collection of alleles for every gene found in the species). Concurrently, laboratory evolution emerged as a new approach to address biological questions and to develop new phenotypic traits, and a large number of laboratory acquired mutations can be found in databases. The availability of this large-scale sequence variation data now allows for a detailed comparison of mutations fixed in natural versus laboratory evolutions. Such comparison shows that laboratory-acquired mutations are rarely found in the wild-type alleleome of *Escherichia coli*. The *E. coli* alleleome is highly conserved as most of the sequence variation is concentrated in about 2% of the coding region. We find that there are typically two alternate amino acids coded for in the variable locations, and switches between the two are found in the data sets. Finally, we find that adaptive laboratory mutations, unlike wild-type mutations, do not utilize the redundancy built into the genetic code: they are less likely to be synonymous and rely on changing a single nucleotide in a codon. However, the uniqueness of mutations fixed in laboratory evolutions bodes well for synthetic biology by revealing novel exploitable sequence space untouched by natural evolution.

## Manuscript

In the late 2000s, DNA sequencing costs dramatically decreased. Inexpensive sequencing led to a steady increase in the number of sequenced genomes in strains of a bacterial species throughout the 2010s ^1–3^. Thus, sequence variation amongst these strains can now be studied at an unprecedented scale. We assembled a collection of 2,661 fully sequenced *E. coli* genomes and defined DNA and amino acid sequence variation amongst them (Fig. 1A). We found that most of the amino acid sequence encoded by a gene (an Open Reading Frame, ORF) was conserved with only a few locations in the ORF where there was notable occurrence of amino acid substitutions. We defined a 3D histogram that showed the amino acid occurrence for every position in a given ORF in this population of *E. coli* strains (Fig. 1B). The simplicity of amino acid substitutions allows us to describe the dominant and minor alleles (variants) on a 3D structure of the protein (Fig. 1C). The particular amino acid with the greatest WT occurrence at each position (dominant amino acid) in a protein is used to define a consensus sequence (WT consensus) for the ORF (Fig. 1D, black). This consensus sequence shows the amino acid positions in the protein that are fully conserved, the positions that are the most variable, and the occurrence and location of significant variants. Since most of the dominant amino acid positions have an occurence close to 100% (i.e. found in all strains that carry that gene), we can prepare a histogram of the dominant amino acid frequencies (0 < c ≤ 1, by normalizing to the number of strains carrying the gene) over all positions to get a simple position-independent view of the sequence variation in the ORF (Fig. 1E). Most ORFs are found in all 2,661 strains, but some ORFs are found in a subset of these strains, see Supplemental Figure 1.

**Figure 1:**
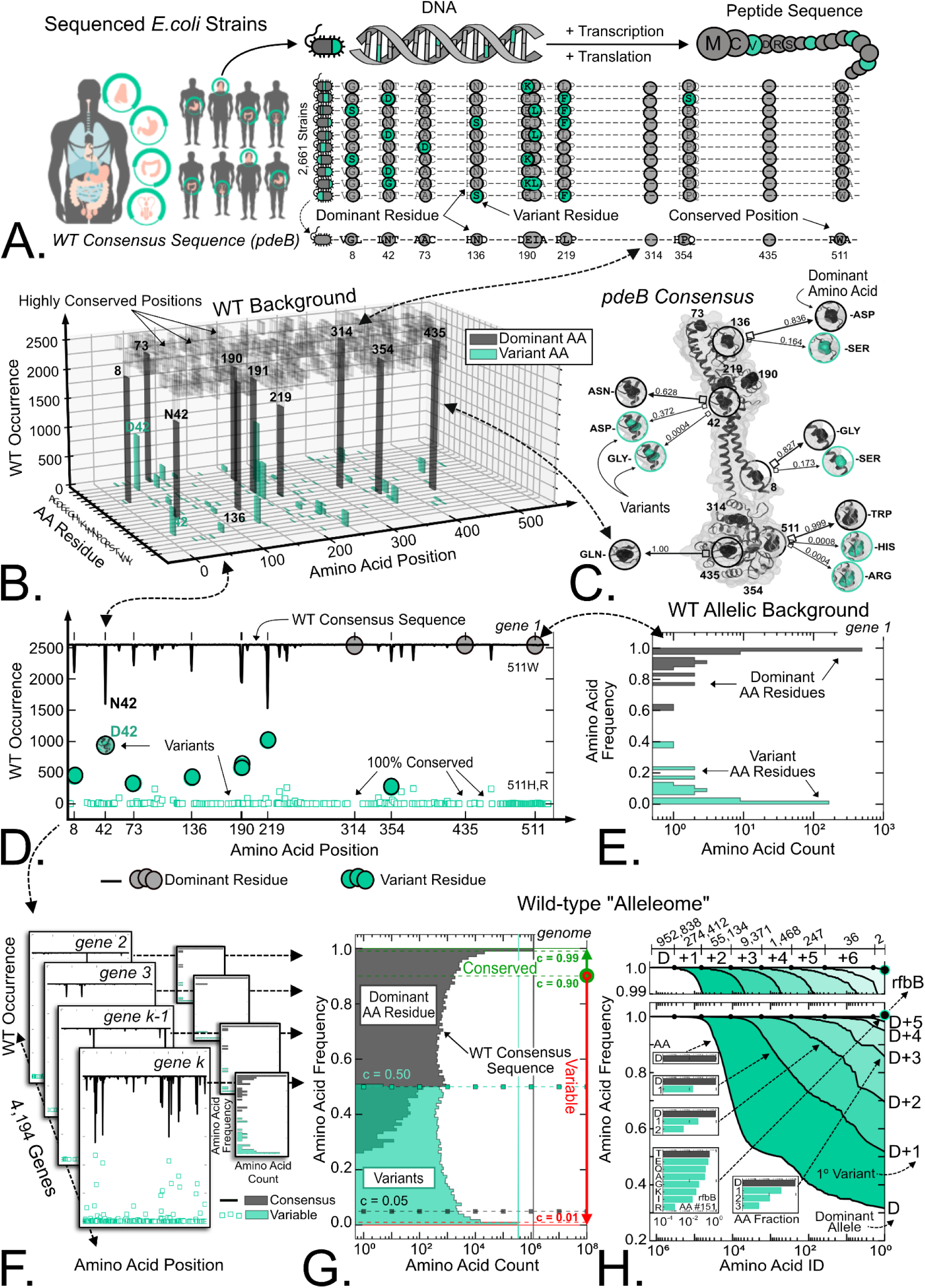
The wild type *E. coli* Alleleome based on whole genome sequences of 2661 wild type strains. **(A)** The sequence variation in 2,661 fully sequenced clinical isolates and commensal strains of *E. coli* was used to determine a wild-type (WT) consensus amino acid sequence for each gene (open reading frame). The phosphodiesterase gene *pdeB* is shown here as an example. Its consensus amino acid sequence–consisting of the amino acid of the highest occurrence (dominant amino acid) in each position–is shown at the bottom of the panel. **(B)** A 3D histogram of the WT occurrence (0<strain count<2661) of every amino acid in every position of *pdeB*. Most amino acid positions have an amino acid of a dominant occurrence, with much lower occurrence of secondary and tertiary amino acids. Few alternative amino acids are found in each of the variable positions. Conserved positions with no variants are shown (black, semi-transparent). This histogram summarizes the alleleome of *pdeB that* reflect non-synonymous mutations in the WT strain collection analyzed. **(C)** Given the simplicity of the occurrence distribution of natural wild-type amino acid substitutions, this variation can be summarized in a 3D rendering of the protein. **(D)** The *WT occurrence* (0 < strain count < 2661) of dominant and variant amino acids, plotted as a function of position in the amino acid sequence. The amino acid sequence of the protein composed of the dominant amino acid in each position is taken to be the consensus amino acid sequence for the ORF (black). Prominent variant amino acids are shown for the most variable positions. **(E)** A histogram of the *dominant amino acid frequency* for all amino acid positions in *pdeB* is shown (gray). Dominant amino acid frequency values (0 < c_position, AA_ ≤ 1) are normalized from the WT occurrence (strain count) values in 1D. The frequency of the secondary and tertiary amino acid variants is also indicated (cyan). **(F)** The individual histograms of WT occurrences in all genes (ORFs) can be generated and combined to show **(G)** the amino acid frequencies in all 1.29 million codon positions that make up the *E. coli Alleleome*. The conserved region of the alleleome is defined as all amino acid positions whose dominant amino acid alleleomic frequency is at least 90% (c ≥ 0.90) (green). Theoretical limits are shown for variant and dominant residues (0 < c ≤ 0.50 and c ≥ 0.05, respectively). By definition, residues present in more than 50% of strains cannot be ‘variants’ while an equal distribution of all 20 amino acids at a specific position would result in a ‘dominant’ residue frequency of 5%. **(H)** Each of the 1.29 million amino acid positions is defined by a dominant amino acid and any additional variants present at that same position. The cumulative amino acid frequency (wild-type diversity) captured by a dominant (D) amino acid and subsequent variant amino acids (+1, +2, …+6) is shown. The amino acid diversity of the *E. coli* alleleome is narrow – 99.1% of positions are defined by three or fewer amino acids. The insert (top) shows the number of amino acid positions for which the wild-type diversity is fully captured by the subset of alleles. There are 952,838 positions for which the wild-type diversity is represented solely by the dominant allele (D) (i.e. there is no additional variant amino acids); 274,412 positions for which the wild-type diversity is represented by a binary choice between the dominant allele and a primary variant (D+1); etc. The average distribution of amino acids for a subset of codons that fully capture the wild-type diversity at a given position is shown (interior inserts) (i.e. for the 55,134 positions with exactly one dominant amino acid and two variant amino acids (D+2), the average distribution of amino acids for the dominant amino acid (D), primary variant (1°) and secondary variant (2°) is 95.1%, 4.6% and 0.36%, respectively). The amino acid distribution is shown in detail for glucose dehydratase protein rfbB at amino acid position 151, the most variable position in the *E. coli* alleleome, where 8 alternate amino acids are present.

These histograms illustrate the *alleleome* for a single ORF, that we can generalize to all the ORFs found in the 2,661 strains. We can combine all histograms (as shown in Fig. 1E) for all these ORFs to generate a combined histogram that reflects the all dominant and variant amino acid frequencies (Fig. 1F). By combining all the histograms for the individual ORFs, we can generate a global histogram that shows the dominant amino acid for every codon in all the ORFs annotated to these sequences (Fig. 1G, gray). The alleleome also provides a global assessment of amino acid variations found in the *E. coli* proteome, reflecting non-synonymous mutations (Fig. 1G, cyan). The alleleome is a global representation of DNA and amino acid sequence variation in a species based on the available whole genome sequences for strains in the species.

These results show that the natural *E. coli* alleleome is highly conserved. To illustrate these results quantitatively, we compiled the full natural sequence variation background of the 2,661 wild-type *E. coli* genome sequences isolated from diverse environments, which corresponds to a collection of 363,131 alleles across 4,194 ORFs containing 1.29 million amino acid positions (Fig. 1G gray, Supp. Fig. 1). We define the conserved region of the alleleome to consist of amino acid positions where the frequency of the dominant amino acid is at least 90% (c ≥ 0.90) (Fig. 1G, green). In this conserved region of the alleleome, there are 952,838 amino acid positions (73.7%) for which there is absolutely no sequence variation (c = 1.0) among the wild-type strains. There are an additional 278,731 positions (21.5%) for which dominant amino acid is found at a rate of 99% (0.99 ≤ c < 1). Finally, there are 36,188 positions (2.8%) where the dominant amino acid frequency is greater than 90% (0.90 ≤ c < 0.99), showing that 1.27 million amino acid positions (98.0%) in the alleleome are ≥ 90% conserved. Apart from the 1.29 million dominant amino acid residues, the diversity of the *E. coli* alleleome is defined by 420,130 unique amino acid variants (Fig. 1G, cyan) distributed across 340,067 positions (Fig. 1H, cyan). By calculating the number of alternate amino acids found in these positions, we determine that the *E. coli* allelome diversity is extremely narrow – 99.1% of all amino acid positions are characterized by 3 or fewer amino acids (i.e. a dominant amino acid and up to two less common variants).

Adaptive laboratory evolution (ALE) is an experimental approach for biological inquiry and a method for developing phenotypic traits^4–9^. Cultures of bacteria are serially passaged in a defined environment until their growth rate does not change notably, and one or more strains from the endpoint are selected for genome sequencing. A collection of the mutations acquired under ALE has been assembled in a publicly available database (ALEdb.org), which has grown exponentially since its inception in 2019^10^. Presently, ALEdb contains 22,045 mutations obtained from 1,864 bacterial isolates from 108 ALE experiments under a wide range of selection pressures. Mutations found in ALEdb are illustrated in Fig. 2A, and they contain both synonymous and non synonymous mutations. ALEdb also provides an additional tens of thousands of unique ALE-acquired mutations that have not yet appeared in peer reviewed publications (Supp. Table 1). Our analysis is based on 45,415 unique mutations found in ALEdb, 25,472 of which are non-synonymous (Supp. Fig. 2).

**Figure 2:**
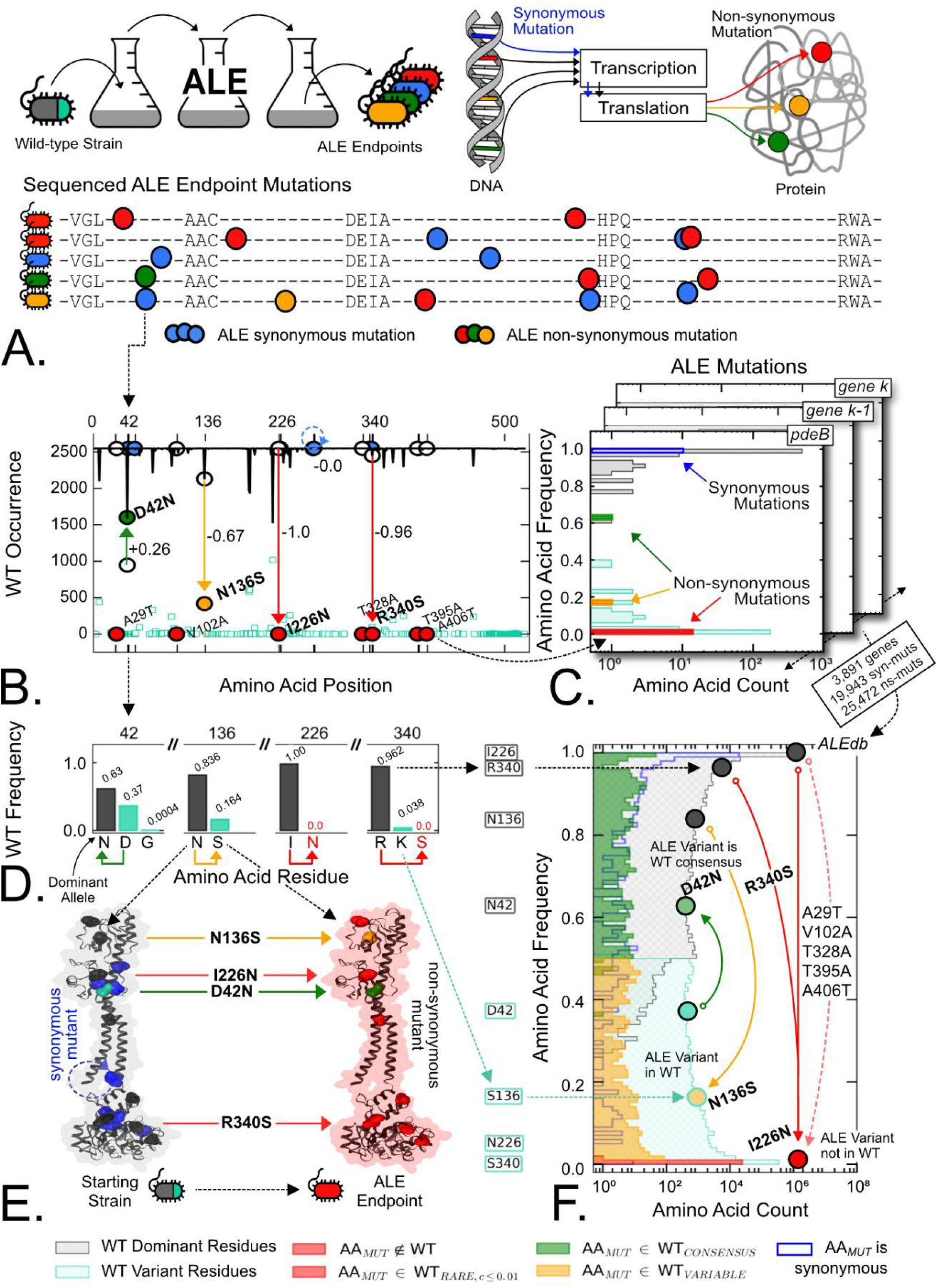
The collection of mutations in *E. coli* strains following adaptive laboratory evolution experiments. **(A)** Adaptive laboratory evolution (ALE) mutations are determined from sequenced genomes of laboratory evolved strains. Synonymous mutations are shown in blue. Non-synonymous mutations resulting in amino acid substitutions i) that are not found or are rarely found (c ≤ 0.01) in the *E. coli alleleome* (red); ii) that are variants in the *E. coli alleleome* (0.01 < c < c_WT consensus AA_) (orange); and iii) that resemble the *E. coli alleleome* consensus amino acid (green) are shown. **(B)** ALE mutations of phosphodiesterase gene, *pdeB*, mapped onto the wild-type consensus amino acid sequence. Arrows connecting the original amino acid residue (white circles) in the starting strain to the observed amino acid substitutions (colored circles) in the ALE endpoint show the change in position-specific alleleomic frequency (normalized WT occurrence) for each ALE mutation. **(C)** A histogram of *WT amino acid frequency* of ALE mutations for all positions, and of dominant and variant amino acids in the wild-type allelic background of *pdeB*. **(D)** The WT frequency of dominant and variant amino acids at four positions of *pdeB* where ALE mutations occur. Arrows indicate ALE non-synonymous amino acid substitutions at each position (e.g. *D42N* reflects an ALE mutation at position 42 resulting from the transition of a starting strain with variant amino acid, *D* (aspartate), into the ALE endpoint with WT dominant amino acid, *N* (asparagine). Note that the alleleome is ‘narrow’ in these positions–there are only two different amino acids found in these positions in most of the 2661 WT strains. **(E)** ALE mutations of *pdeB* are shown on a rendering of its 3D protein structure. Mutated residues in the starting strain are dominant amino acids (black) or variants (cyan) in the *E. coli* alleleome. Synonymous mutations are also shown in blue. Mutated residues in the ALE endpoint are highlighted using the color scheme in Fig. 2A. **(F)** The ‘movement’ of ALE mutations against the histogram of dominant amino acid frequencies of the 1.29 million codon positions that make up the wild-type *E. coli Alleleome* (gray / cyan), that is reproduced from Fig. 1G. The amino acid frequencies of all 25,472 non-synonymous ALE mutations across 3,891 genes found in ALEdb (the green, orange and red color schema is defined in Fig. 2A) are shown. The WT alleleome is used to contextualize specific mutations in *pdeB* leaving (from gray) or returning (from cyan) to the wild-type consensus amino acid. Synonymous ALE mutations are shown in blue.

We can assess the genetic differences between laboratory (‘synthetic’) evolution and natural evolution by comparing the wild-type alleleome and mutations found in ALEdb. We begin by looking at the mutations fixed during ALE in the *pdeB* gene and display them on the WT amino acid occurrence diagram (i.e., Fig. 1D). The result, graphed in Fig. 2B, reveals three types of non-synonymous mutations: first, there are 7 distinct mutations occurring in conserved positions where the amino acid substitution falls outside of the WT alleleome (shown in red); second, an amino acid substitution resulting in a switch from an amino acid of lower occurrence (a WT variant) to the amino acid of dominant occurrence (D42N, shown in green) that can be thought of being a ’revertant’ to consensus; and third, there is an amino acid substitution from a dominant occurrence to one of a lower occurrence (N136S, shown in orange). Thus, ALE can provide a selection pressure that selects for a non-consensus amino acid. The frequency and type of amino acids where substitution takes place are detailed in Fig. 2D, and with such few mutations in *pdeB*, they can be viewed on the 3D protein structure (Fig. 2E). Thus, 7 of the 9 non-synonymous ALE mutations found in *pdeB* fall outside of the WT alleleome, while the remaining two are found in two variable positions in the protein.

As for the WT alleleome (Fig. 1 F&G), mutational data from ALE can be scaled up from one ORF to all the 3,891 ORFs with ALE-acquired mutations. The results, shown in Figure 2C & F, give us a global view of all the acquired ALE mutations relative to the WT alleleome. To assess the mutations in ALEdb, we identify two features of each mutation: first, the frequency with which the amino acid in the starting ALE strain (i.e., MG1655) is found in the WT alleleome (i.e. is the original residue identical to the WT consensus residue or a WT variant residue) and second, the frequency with which the amino acid substitution in the ALE endpoint is found in the WT alleleome (i.e. is the mutation is novel, rare, a WT variant, or a reversion to the WT consensus). Of the 45,415 observed mutations in ALEdb, we find that the majority of synonymous and non-synonymous mutations (98.7% and 96.0%, respectively) occur in conserved consensus positions (c ≥ 0.90) of the WT alleleome (Fig. 2F, blue and Fig. 3C, black). Analyzing the non-synonymous amino acid substitutions acquired through ALE, we find that the majority (96.7%) of the resulting amino acid substitutions are rarely (c ≤ 0.01) found in the WT sequence variation (Fig. 2F & Fig. 3A & Fig. 3B, red). In fact, 88.5% of all ALE acquired mutations are not found in the WT alleleome (Fig. 2F & Fig. 3A & Fig. 3B (‘is Novel), red).

**Figure 3:**
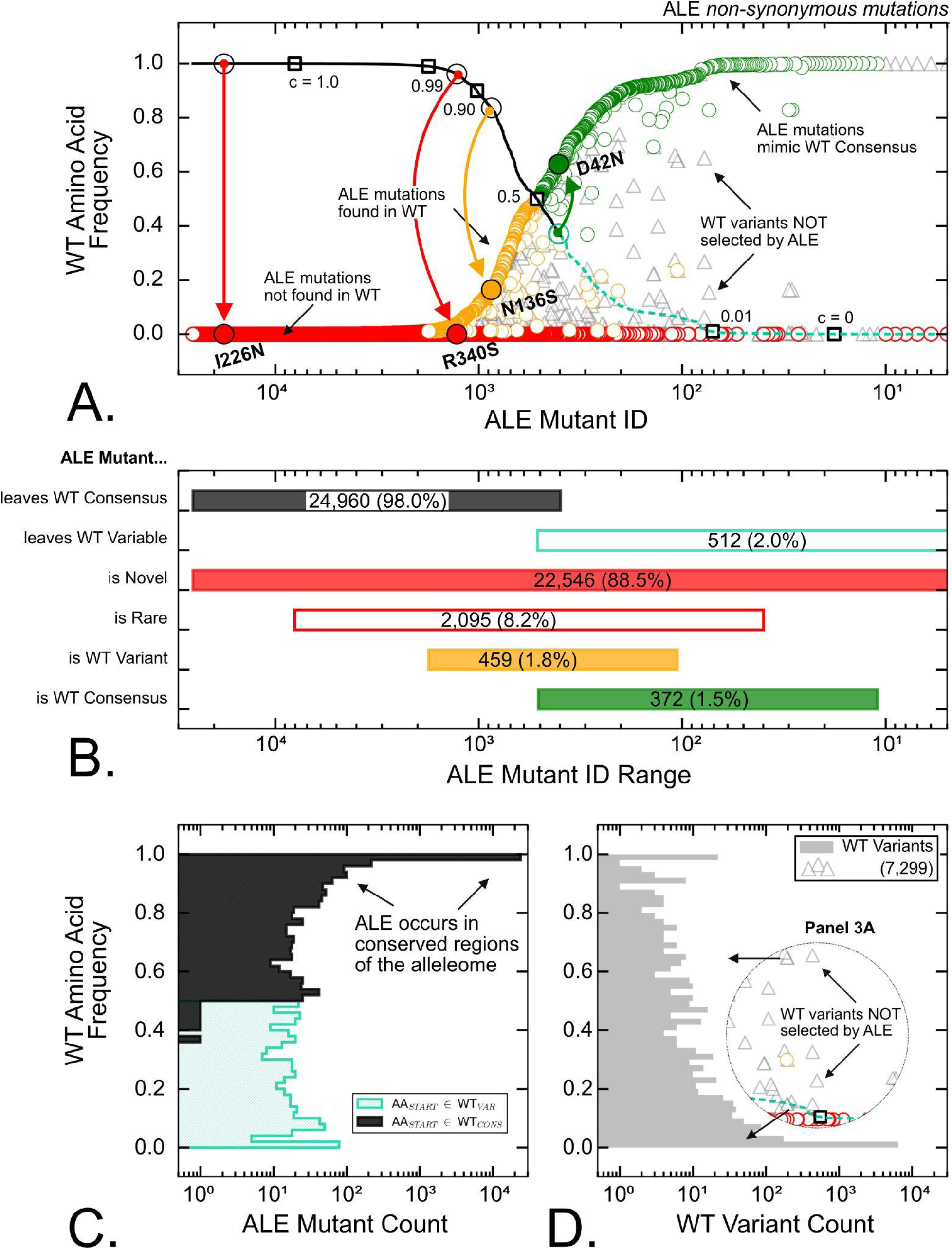
Detailing the set of mutations observed during adaptive laboratory evolution. **(A)** 25,472 non-synonymous ALE mutations were rank-ordered by the wild-type amino acid frequency of the original amino acid in the ALE starting strain. ALE non-synonymous are found leaving the consensus allele (*black line*) or leaving variant alleles (*dotted cyan line*). The frequency of the resulting amino acid substitution in the ALE endpoint is plotted (colored circles) according to the color schema in Figure 2A. Naturally occurring WT variants that are NOT selected for by ALE are also shown (gray triangles). ALE mutations in phosphodiesterase gene (*pdeB)* from Figure 2 are highlighted. **(B)** Non-synonymous ALE mutants are grouped by the amino acid in the starting strain (gray / cyan) and by the amino acid substitution in the ALE endpoint (red / orange / green). The range of ALE mutant IDs and raw number of ALE mutations in each group are shown. **(C)** A histogram of the wild-type alleleomic frequency of the starting strain amino acids is shown. Most non-synonymous ALE mutations are found leaving conserved positions of the WT consensus sequence. **(D)** A histogram of the wild-type alleleomic frequency of variants that are NOT selected for by ALE and do not resemble the starting strain. Many rare WT variants are yet to be selected for by ALE.

By studying the relationship between the starting WT and ALE amino acid frequencies on the large scale that the available data sets offer, intricate characteristics of non-synonymous ALE mutations are revealed. Non-synonymous ALE mutations are rank-ordered by the wild-type frequency of their original amino acid (pre-mutation) (Fig. 3A). We can identify all mutations from residues originally belonging to the WT consensus (Fig. 3A & Fig. 3B & Fig. 3C, black) or from residues that are themselves WT variants (Fig. 3A & Fig. 3B & Fig. 3C, cyan). Mutations that depart from the WT consensus can result in mutations that are novel (red), rare (red/white), or are WT variants in the natural alleleome (orange) (Fig. 3A & Fig. 3B). We note that the latter case represents ALE conditions that select for a WT variant. Mutated residues that were initially WT variants in the starting strain can result in all the aforementioned types of mutations, but can also return to the wild-type consensus residue (Fig 3A & Fig 3B, green), i.e. a reversion to consensus. Since the majority of ALE mutations occur in highly-conserved consensus regions of the natural alleleome and result in amino acid residues that are not found in the WT sequence variation, it is likely that laboratory and natural selection pressures are largely disjoint. This divergence thus suggests that serial passaging may subject laboratory strains to significantly different evolutionary pressures than those experienced by wild-type strains.

The existence of ALE mutations which result in WT variants of the natural alleleome (459, orange) and those that revert to the WT consensus (372, green) suggest that a small subset of ALE selection pressures overlap with the wild-type selection in natural strains (Fig. 3B). These observations may guide a new generation of future ALE experimental design. For instance, ALE mutants that revert to the WT consensus in one experiment can be subjected to a range of selection pressures until the mutant returns back to the original amino acid in the first experiment. Identifying these selection pressures that can wobble specific amino acids between wild-type consensus residues and variants may offer insights into divergence in evolution between natural ‘wild-type’ strains and laboratory ‘wild-type’ strains.

Additionally, by identifying 7,299 wild-type variants (Fig. 3A & Fig. 3D, gray triangles) in the genomic positions where ALE mutations are found, experiments can be designed to identify new selection pressures that select for these currently untouched natural variants, strengthening our understanding of the evolutionary trajectories of natural strains. Given the focused protein structure consequences of single amino acid substitutions, wobbling specific amino acid positions using ALE selection pressures can give us a new class of structure-function mutations, where the function is at the physiological level, not the protein level. These mutations thus have a systems biology relevance.

Both natural evolution and laboratory evolution are subject to the constraints imposed by the redundancy of the genetic code itself. Not all amino acid substitutions are achievable by a single point mutation. In many cases, to achieve a particular amino acid substitution requires two or even three point mutations. As an example, we show the mutation trajectory required for a UUU_0_→GGG_3_ codon change in Figure 4A, which is one of eight possible codon-changes that result in the phenylalanine → glycine amino acid substitution. In this trajectory, the UUU codon must acquire three sequential point mutations before reaching the final GGG codon. Intermediates along the trajectory UUU_0_→GGG_3_ are denoted by the number of point mutations (‘hops’) they have acquired.

**Figure 4:**
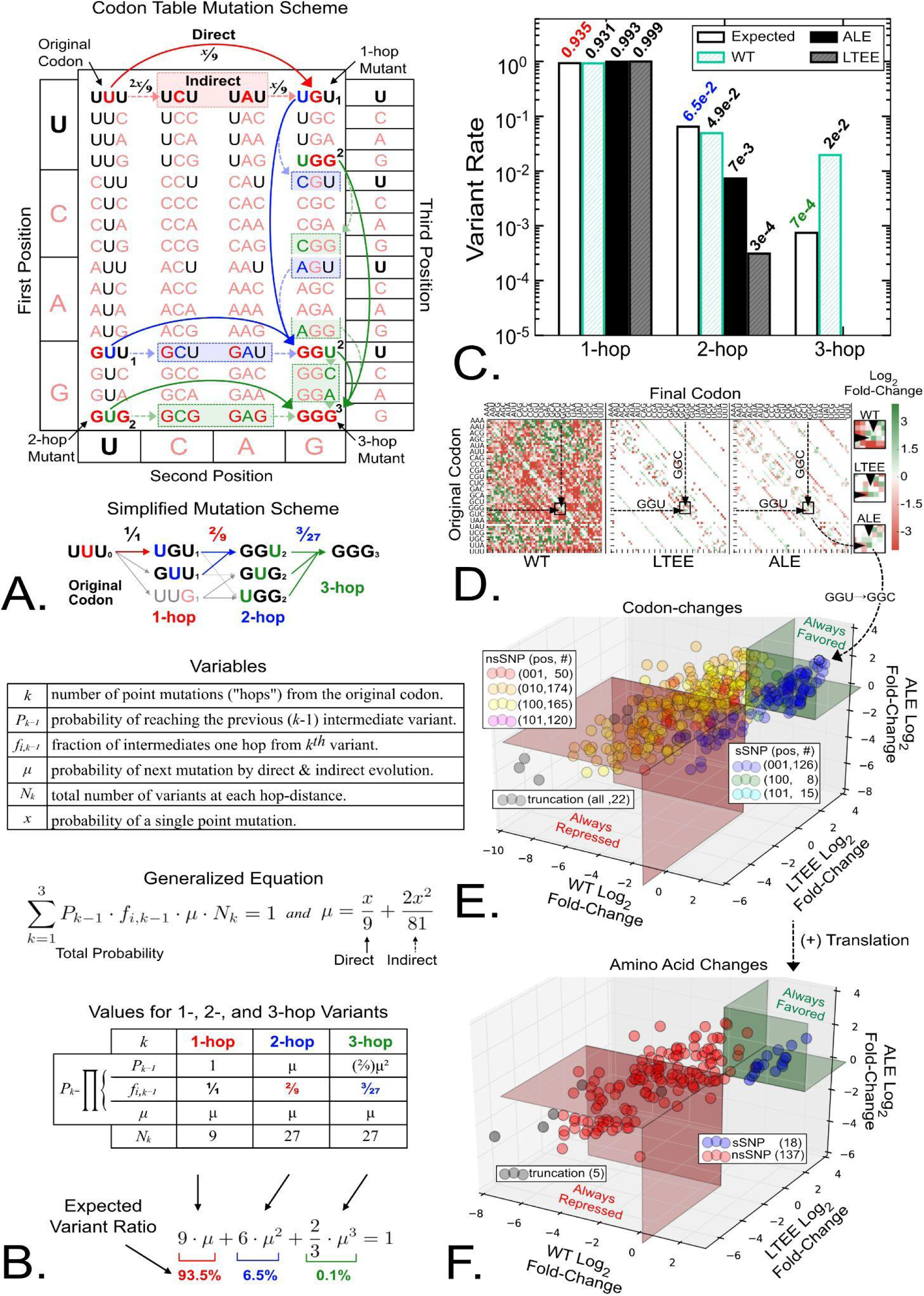
Observed mutant ratios in both laboratory and natural evolution are constrained by fundamental mutation trajectories. **(A)** In many cases, two or even three point mutations at the codon-level are needed to achieve a particular amino acid substitution. The UUU to GGG codon-change is one of eight phenylalanine → glycine amino acid substitutions. The mutation trajectory following path UUU→UGU^1^→GGU^2^→GGG^3^ is shown in detail. For any codon-change, the new codon is referred to as an *k-hop* mutant, where *k* is the number of different nucleotides between the new codon and the original codon. Subscripts 1, 2 and 3 denote the minimum number of hops required to reach the mutant. Any codon-change that deviates from the previous codon by exactly one nucleotide (1-hop) can be achieved by a single point mutation (correct nucleotide and position) (direct evolution), or by two point mutations (the first places the wrong nucleotide at the correct position) (indirect evolution). A simplified mutation schema for UUU→GGG ^3^ and its intermediates is shown. **(B)** Method for calculating the expected distribution of mutations 1-hop, 2-hops and 3-hops away from any given starting codon. In short, the probability of reaching a specific *k*-hop mutant (P_*k*_) is the product of: i) the probability of reaching the previous mutant (P_*k-1*_); ii) the fraction of previous intermediates that are can be reached one-point mutation prior to the *k*^*th*^ mutant (*f*_*i,k-1*_); and iii) the probability of reaching the desired *k*-hop mutant through direct & indirect evolutionary trajectory (*μ*). A single point mutation can result in one of nine unique codon-changes, such that each subsequent specific 1-hop codon-change can be reached directly 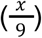 or indirectly 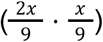, where x is the probability of a point mutation. The sum of the probabilities of reaching all (1-, 2- and 3-hop) mutants for each starting codon must be equal to one. The appropriate values for all 1-3 hop mutants and expected distribution are shown. **(C)** The expected (i.e., calculated) and observed rates (WT, ALE, LTEE) for all 1, 2, and 3-hop mutants are shown. **(D)** The log_2_ fold-change (from the expected value) in the rate of observed specific codon-changes for the wild-type *E. coli* variants, ALE mutants and LTEE mutants. **(E)** The log_2_ fold-change values for specific codon-changes found in all of the datasets (WT *E. coli* variants, ALE, and LTEE mutants) is graphed in 3D to identify specific codon-changes that are always enriched in the observed data (green) or always under-represented in the observed data (red). Codon-changes are colored by the nucleotide position that is mutated and their protein phenotype (synonymous, non-synonymous, truncation) (i.e. 50 non-synonymous 1-hop mutations where the third nucleotide in the codon is mutated are shown in red, nsSNP). **(F)** Specific codon-changes are grouped by the amino acid residue they code for. The expected distributions of synonymous (blue) and non-synonymous amino acid substitutions (red) are calculated and deviations (log_2_ fold-change) from the expected expected value can be determined as in Figure 4E.

To determine the consequences of the genetic code’s constraint on natural variation and laboratory evolution, we develop a simple mathematical model to quantitatively predict the distribution of codon-changes and subsequent amino acid substitutions achievable from a given codon (Fig. 4B). The computation is governed by two simple principles. First, any specific “1-hop” codon-change can be reached by a single point mutation at the correct position and with the correct nucleotide (direct evolution) (i.e., UUU→UGU), or by two mutations, the first being a substitution of the wrong nucleotide in the correct position (indirect evolution) (i.e., UUU→U(C or A)U→UGU). Second, the probability of achieving a multiple-hop codon-change is dependent on the probability of passing through all suitable intermediates along the mutation trajectory (i.e., P_UUU→UGG_ = Σ P_UUU→*i*_ · P_*i*→UGG_ where *i* ∈ {UGU,UUG}). The complete computation of the probabilities is shown in Figure 4B and they predict 1-, 2-, and 3-hop codon-changes will comprise 93.5%, 6.5%, 0.1% of the variants for any given codon, respectively.

To identify differences in mutations found in natural evolution (WT alleleome), adaptive laboratory evolution (ALE) and long-term evolution experiment (of multi-decade duration) (LTEE) pioneered by Lenski^12,13^, we add an additional dataset of 10,572 unique mutations acquired in ORFs through LTEE for subsequent analysis (https://barricklab.org/shiny/LTEE-Ecoli/). The LTEE mutations can be analyzed as the ALE mutations (Supplemental Fig. 3) that yields similar results as those for ALE in Figs. 2 and. 3.

At the genome-level, we find that our simple model accurately reproduces the distribution of 1-, 2-, and 3-hop codon-changes for natural sequence variation and laboratory evolution (Fig. 4C). The experimental data shows that codon-changes that require multi-hop mutations are much more frequent in the WT alleleome than in adaptive laboratory evolutions and long-term laboratory evolutions. In fact, none of the 55,987 laboratory mutations (ALE + LTEE) require 3-hops. Using the computational model, we can determine whether observed individual codon-changes are enriched in either the wild-type alleleome, ALE, or LTEE mutations (Fig. 4D). Enrichment of codon-changes found in all three datasets is shown (Fig. 4E). Redundancy in the genetic code allows for multiple codons to code for the same amino acid, therefore all codon-changes resulting in the same amino acid substitution can be grouped together to show the enrichment of specific amino acid substitutions common to all three datasets (Fig. 4F). Further analysis (Supp. Fig. 4-7), shows that the occurrence of synonymous mutations is likelier in the WT alleleome than in laboratory evolution. By analyzing the intricacies of the genetic code we thus discover two different properties of WT vs laboratory evolutions: i) laboratory evolutions utilize fewer multi-hop codon-changes and ii) are less likely to fix synonymous amino acid substitutions.

Inexpensive DNA sequencing has allowed us to define natural sequence variation amongst wild-type strains in the form of the *alleleome*. We find a large fraction of the alleleome to be conserved and have a dominant occurrence of a particular amino acid (98% of positions have a dominant amino acid present in ≥90% of strains). We find that about 2% of the 1.29 million codons in *E. coli* to be variable but typically only have a single amino acid of secondary dominance. Thus, the WT alleleome of *E. coli* is relatively ‘narrow’ (Fig. 1H).

Inexpensive DNA sequencing has also enabled us to resequence a large number of clones from laboratory evolutions to find causal mutations relative to a specific selection pressure. This first large-scale comparison of acquired mutations during experimental evolution with the WT alleleome shows that they are largely divergent, suggesting that the selection pressures that have been used in experimental evolutions to date do not reflect those experienced during natural selection. Remarkably, there is a subset of mutations observed in ALE which suggests that well-defined selection pressures can switch between the dominant and secondary dominant amino acids, and vice versa.

The uniqueness of the vast majority of experimental evolution mutations is advantageous from a synthetic biology point of view. ALE has been shown to be useful to generate desirable bioprocess phenotypes, such as those exhibiting increased tolerance for end-product toxicity and improved substrate readiness^6^. This cumulative experience shows that there is useful sequence space to be explored for biological design purposes, and it may be non-obvious and thus advantageous from a discovery standpoint.

## Author contributions

E.C. and B.O.P. designed the study. P.P. contributed to the acquisition of ALE mutation data. J.M. contributed to the acquisition of WT sequence data. E.C. performed data curation, technical analyses and modeling. E.C. and B.O.P. drafted the paper. All authors helped draft and edit the final paper.

## Acknowledgements

We would like to thank Adam Feist for help with curating and managing mutations in ALEdb, Siddharth Chauhan and Daniel Zielinski for helpful technical discussions, and Marc Abrams for assistance with manuscript editing. This work was funded by the Novo Nordisk Foundation (Grant Number NNF20CC0035580) and by the National Institutes of Health (Grant R01 GM057089).

