## Supplemental Figures 1-7 for "Laboratory-acquired mutations fall outside the wild-type alleleome of *Escherichia coli*"

Edward Catoiu, Patrick Phaneuf, Jonathan Monk, Bernhard Palsson
Department of Bioengineering, UCSD, La Jolla CA 92101

### Supplementary Figures

#### Supp. Fig. 1: Strain, gene and nucleic acid allele counts from sequenced natural *E.coli* strains of the WT alleleome.

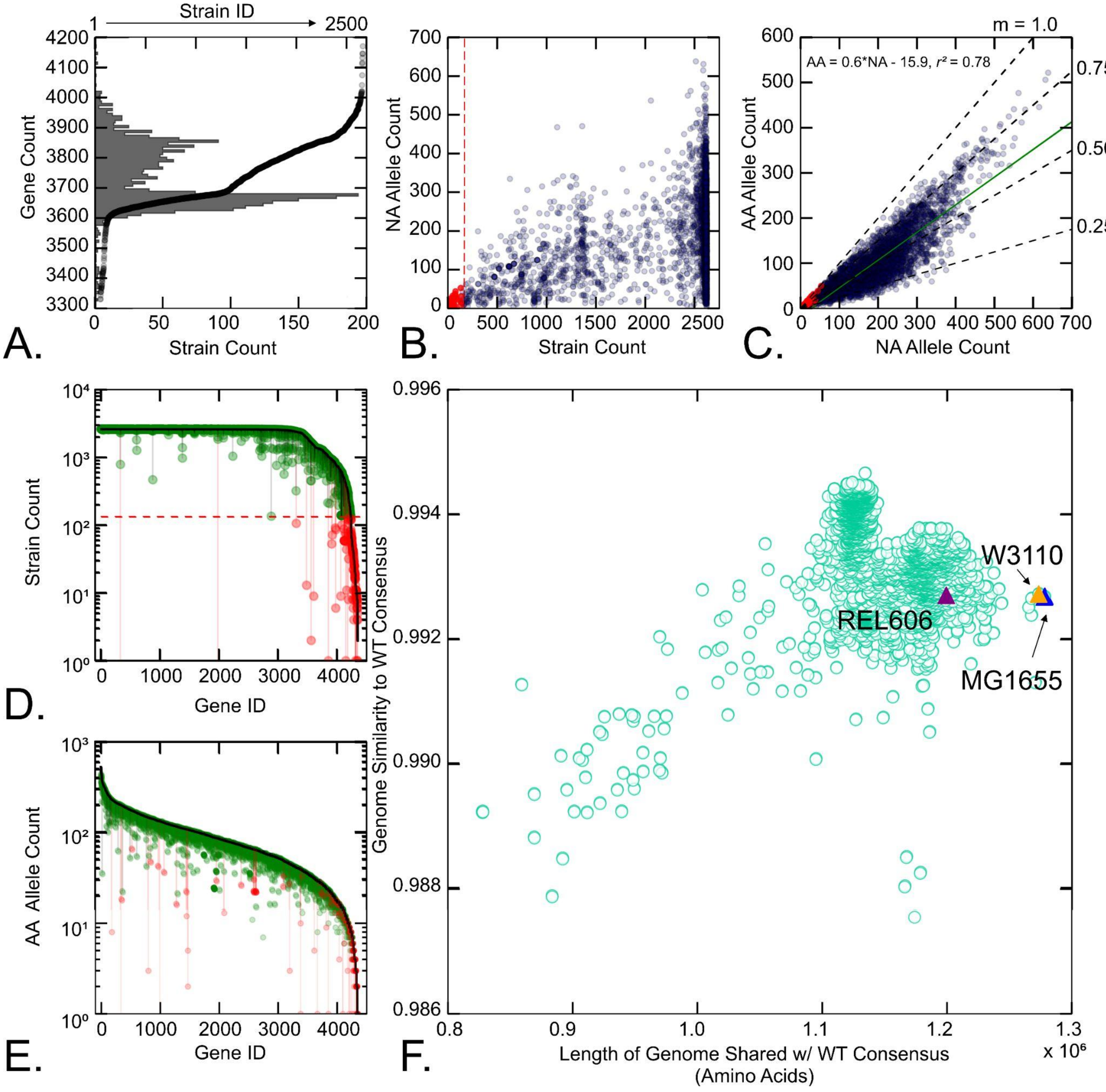


(A) The number of genes in each of the 2,661 sequenced wild-type *E. coli* strains (black). A histogram of the number of genes in each of the wild-type *E. coli* strains (gray).

(B) The distribution of 778,250 distinct nucleic acid alleles for 4,349 genes Open Reading Frames (ORFs) across 2,661 wild-type *E.coli* strains. Genes present in less than 5% of strains are removed from subsequent analysis (red).

(C) Since the genetic code is redundant, multiple nucleic acid allele sequences can be translated into the same amino acid allele. When analyzed at the amino acid level, the natural variation space is reduced by the effect of synonymous mutations. This effect is shown here. The number of nucleic acid alleles in Panel B is reduced to 410,007 unique amino acid alleles.

(D) QC/QA analysis of amino acid alleles was performed. Alleles were eliminated from the population if it results in a peptide sequence : i) truncated by more than 20% or ii) more than two standard deviations shorter than the mean peptide sequence length among all alleles.The number of strains for a given gene represented by all its amino acid alleles alleles (pre- and post-QC/QA) is shown. Genes no longer found in more than 5% of strains (133 strains) are removed from subsequent analysis (red). The strain count is used as an upper bound when normalizing WT occurrences to obtain alleleomic frequencies for each gene in Figures 1-3.

(E) Number of amino acid alleles found for each gene pre- and post-QC/QA. The *WT alleleome* is composed of a total of 363,131 unique QC/QA'd amino acid alleles distributed across 4,194 ORFS that exist in more than 5% (> 133) of the strains (green). Sequence alignment of these alleles is used to determine the WT consensus sequence for each gene (Fig. 1).

(F) Following the determination of the WT consensus sequence, the genome similarity to the consensus can be calculated for each of the wild-type strains. *E. coli* MG1655 shares 4,109 ORFs with the WT consensus and has a sequence similarity of 99.268%. *E. coli* REL606 shares 3,809 ORFs with the WT consensus and has a sequence similarity of 99.271%.

#### Supp. Fig. 2: Curation of public and private data in ALEdb, and LTEE mutations.

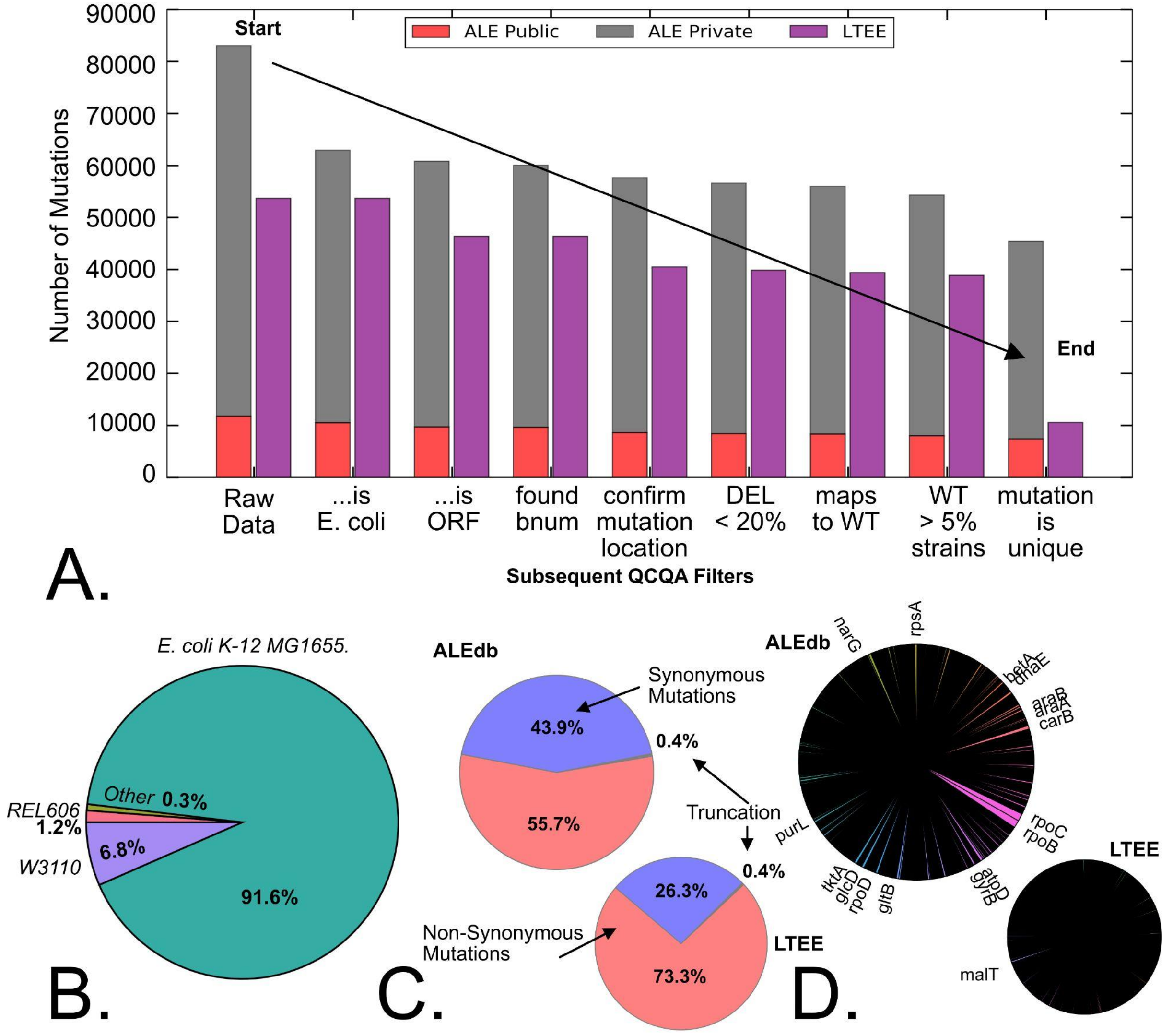


(**A**) QC/QA filters are applied to private and public mutation data from ALEdb, as well as the LTEE data. From left to right: (i) raw data, (ii) experiments in *E. coli* strains with sequenced reference genomes available, (iii) mutations found in genes (Open Reading Frames, ORFs), (iv) mutations where Blattner numbers (Bnums) can be identified, (v) genome position and amino acid position and residue from reference genome sequence matches the mutation annotation provided, (vi) if the mutation is a deletion, the resulting truncation can be no larger than 20% of the full gene length, (vii) the gene and amino acid position is found in the WT alleleome and (viii) is present in at least 5% (133 of 2,661) of strains, (ix) the mutation is only used once (unique mutations). This study gathered 45,415 QC/QA'd unique mutations from the public and private

ALE mutation data (7,392 and 38,023, respectively), and 10,572 QCQA'd mutations from long term evolution experiments hosted at https://barricklab.org/shiny/LTEE-Ecoli/.
(**B**) The majority of ALE mutations come from experiments involving *E. coli* MG1655.
(**C**) The mutation dataset includes mutations that result in synonymous and non-synonymous amino acid substitutions, as well as early protein truncations.
(**D**) The most commonly mutated genes in ALEdb and LTEE data.

### Supp. Fig. 3: Movement of LTEE mutations against the alleleome background.

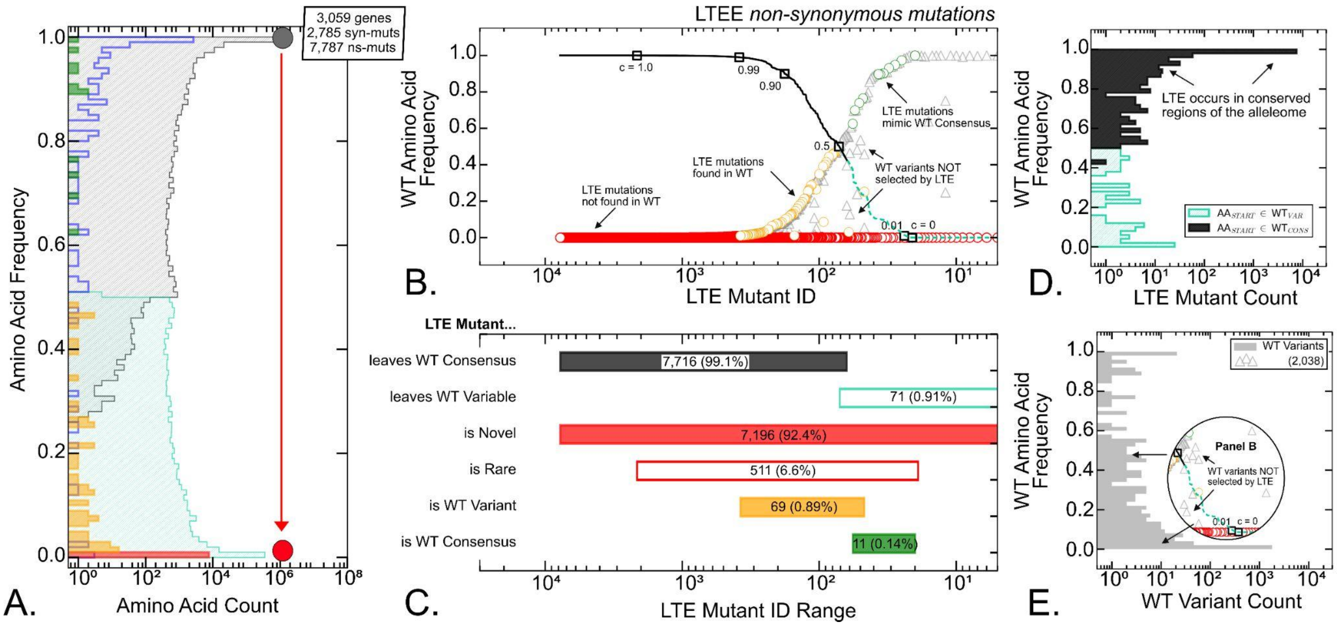


(**A**) The 'movement' of LTEE mutations against the histogram of dominant amino acid frequencies of the 1.29 million codon positions that make up the WT *E. coli Alleleome,* that is reproduced from Fig 1G. The WT data is used to contextualize the amino acid frequencies of all 7,787 non-synonymous mutations across 3,059 genes found in LTEE (the green, orange and red colors are as defined in Panel C). Synonymous LTEE mutations are also shown in blue.
(**B**) 7,787 non-synonymous LTEE mutations were rank-ordered by the wild-type amino acid frequency of the original amino acid in the reference strain (REL606). LTEE non-synonymous amino acid substitutions are found leaving the WT alleleome consensus sequence (*black line*) or leaving secondary variants (*dotted cyan line*). The frequency of the resulting amino acid substitution in the LTEE strain is plotted (colored circles) according to the color schema in Panel C. Naturally occurring secondary WT variants that are NOT found represented in LTEE mutations are also shown (gray triangles).
(**C**) Non-synonymous LTEE mutants are grouped by the amino acid in the reference strain (REL606) and by the amino acid substitution. The range of LTEE mutant IDs and raw number of LTEE mutations in each group are shown.
(**D**) A histogram of the wild-type alleleomic frequency of the reference strain amino acids is shown. Most non-synonymous LTEE mutations are found leaving the WT consensus sequence.
(**E**) A histogram of the wild-type alleleomic frequency of secondary variants that are NOT found in LTEE mutations and do not resemble the reference strain (REL606).

#### Supp. Fig. 4: Log$_2$-Fold Enrichment of codon-changes in WT, ALE, and LTEE compared to expected values (computed from model in Figure 4B).

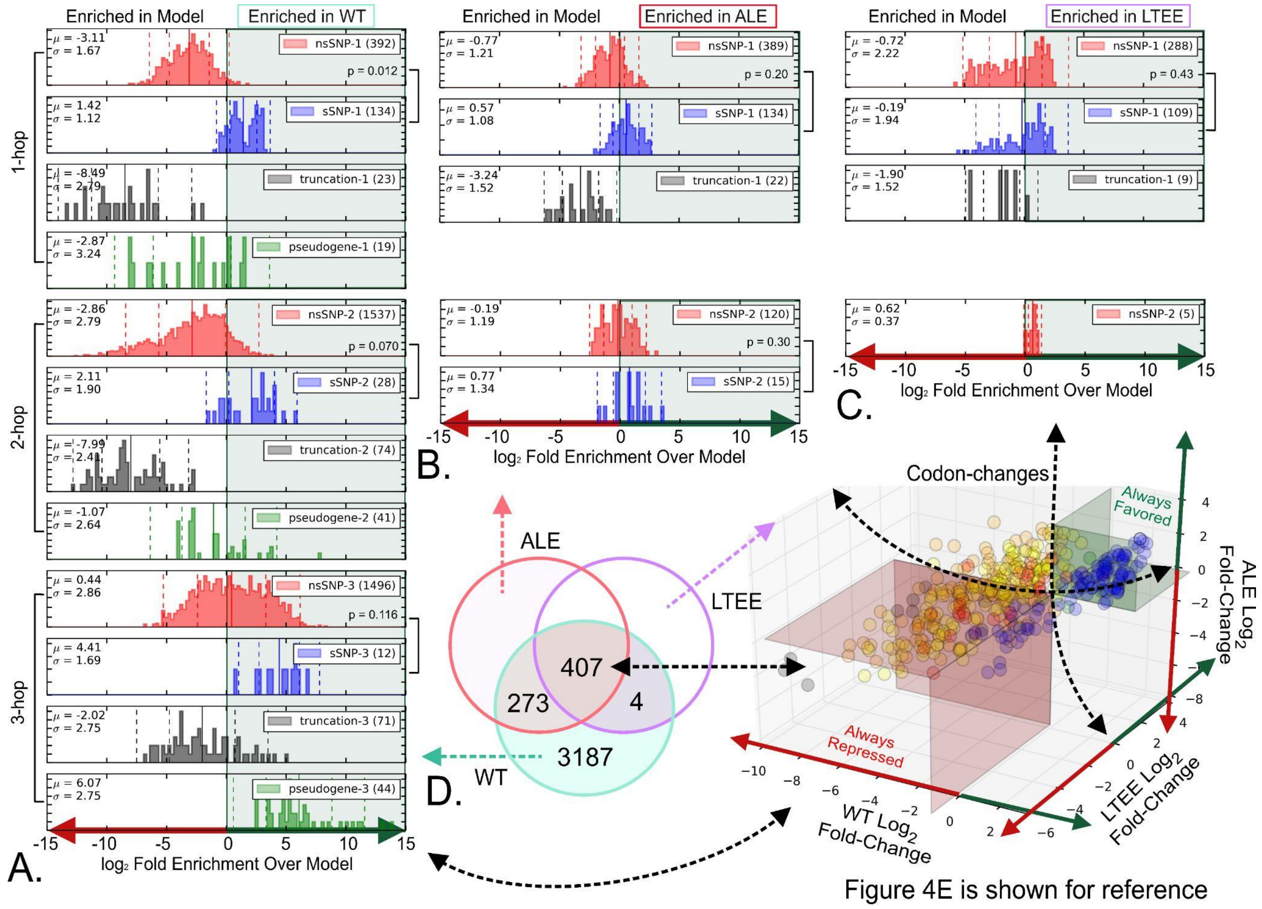


**(A)** Enrichment (log$_2$ fold-change) over model expectations for all 3,871 codon-changes found in WT variants. 1-hop, 2-hop and 3-hop codon-changes are shown separately to normalized differences across multiple-hop trajectories. Mean value and standard deviation are shown for each distribution. At the 1-hop level, WT mutations show a significant enrichment in synonymous mutations ($p = 0.012$).

**(B)** Enrichment (log$_2$ fold-change) over model expectations for all 680 codon-changes in ALE. 1-hop and 2-hop codon-changes are shown separately to normalized differences across multiple-hop trajectories. Mean value and standard deviation are shown for each distribution. The enrichment of synonymous mutations found in WT data is largely attenuated.

**(C)** Enrichment (log$_2$ fold-change) over model expectations for all 411 codon-changes in LTEE. 1-hop and 2-hop codon-changes are shown separately to normalized differences across multiple-hop trajectories. Mean value and standard deviation are shown for each distribution. The enrichment of synonymous mutations found in WT data is largely attenuated.

**(D)** Distribution of codon-changes among the WT, ALE, and LTEE datasets. WT, ALE, and LTEE figures (Panels A-C) were created using 3,871, 680, and 411 unique codon-changes, respectively. The 407 codon-changes common to all three datasets are shown in Figure 4E.

### Supp. Fig. 5: Pairwise comparisons of WT, ALE, and LTEE codon-changes show relative differences in enrichment.

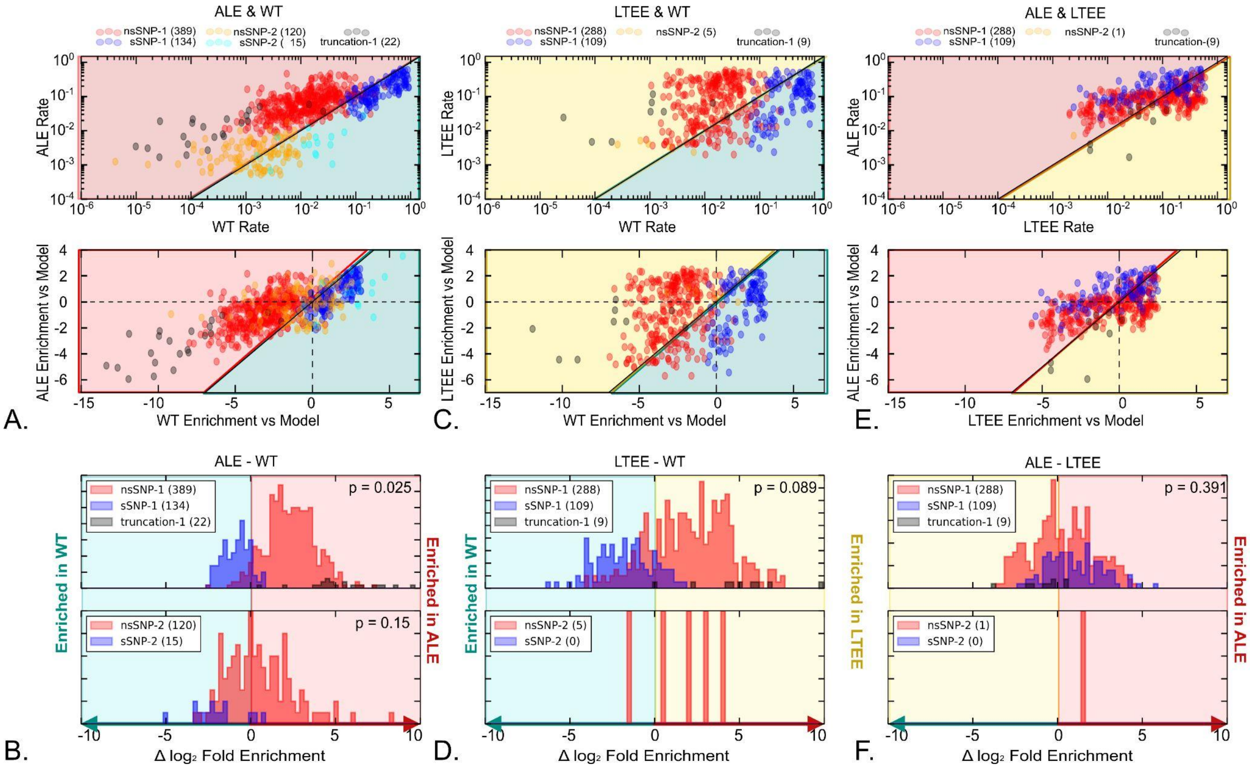


**(A)** Rate of occurrence 680 codon-changes found in both ALE and WT data (top). The occurrence rates are normalized ($log_2$-fold enrichment) using the expected value (bottom). Synonymous 1-hop (blue) and 2-hop (cyan) mutations are shown. Non-synonymous 1-hop (red) and 2-hop (orange) mutations are shown. Truncations are shown (gray).

**(B)** For each codon-change, the relative change in enrichment is calculated. The distribution of this relative change is shown for 1-hop mutations (top) and 2-hop mutations (bottom). When compared to ALE, WT mutations are more likely to fix synonymous 1-hop mutations ($p = 0.025$).

**(C)** Rate of occurrence 411 codon-changes found in both LTEE and WT data (top). The occurrence rates are normalized ($log_2$-fold enrichment) using the expected value (bottom). Synonymous 1-hop (blue) mutations, non-synonymous 1-hop (red) and 2-hop (orange) mutations are shown. Truncations are shown (gray).

**(D)** For each codon-change, the relative change in enrichment is calculated. The distribution of this relative change is shown for 1-hop mutations (top) and 2-hop mutations (bottom). WT mutations are more likely to fix synonymous 1-hop mutations relative to LTEE.

**(E)** Rate of occurrence 407 codon-changes found in both ALE and LTEE data (top). The occurrence rates are normalized ($log_2$-fold enrichment) using the expected value (bottom). Synonymous 1-hop (blue) mutations, non-synonymous 1-hop (red) and 2-hop (orange) mutations are shown. Truncations are shown (gray).

**(F)** For each codon-change, the relative change in enrichment is calculated. The distribution of this relative change is shown for 1-hop mutations (top) and 2-hop mutations (bottom).

#### Supp. Fig. 6: Log$_2$-Fold Enrichment of amino acid substitutions in WT, ALE, and LTEE compared to expected values (model).

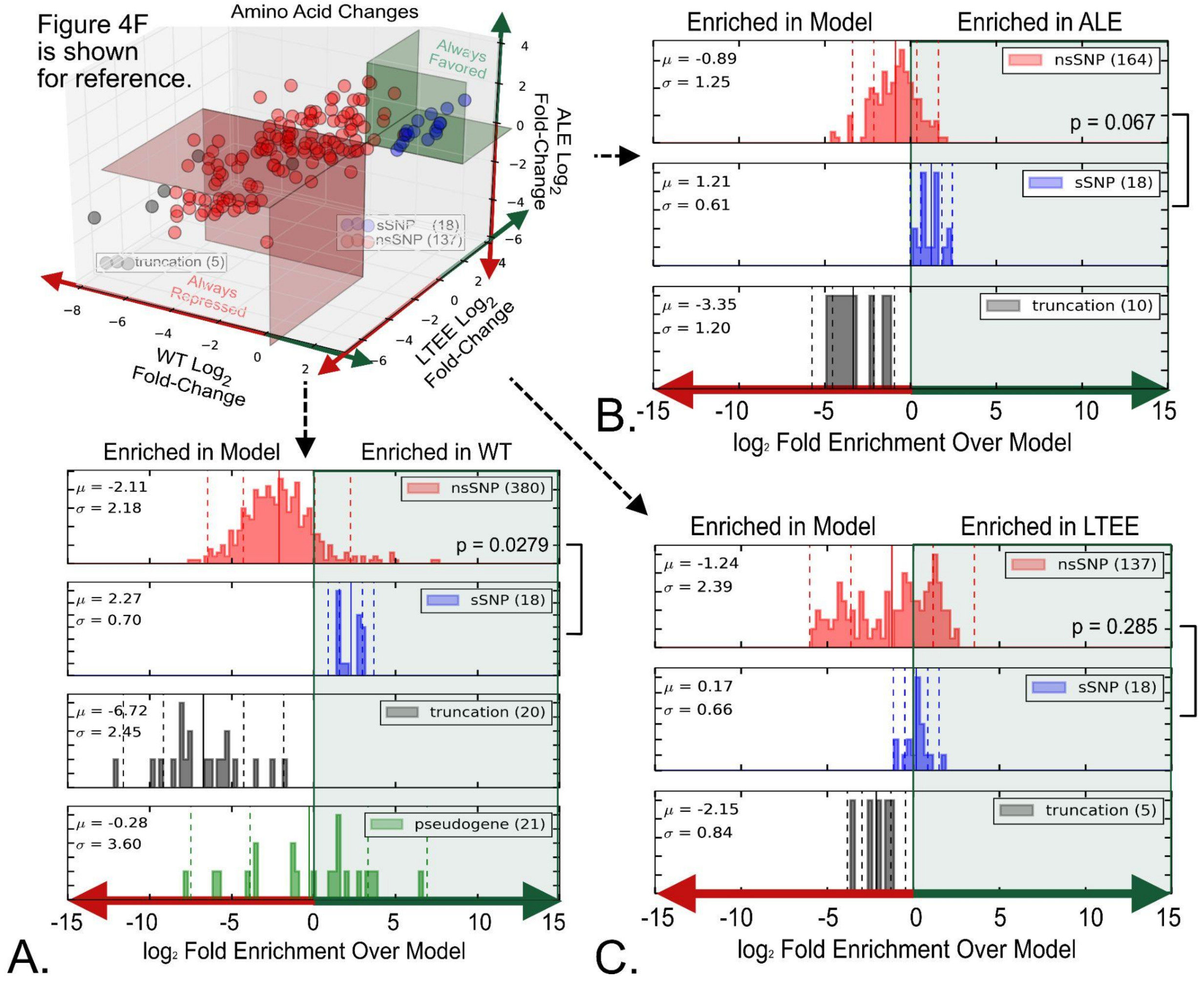


**(A)** Enrichment (log$_2$ fold-change) over model expectations for all 439 amino acid substitutions found in WT variants. Mean value and standard deviation are shown for each distribution. WT mutations show a significant enrichment in synonymous mutations ($p = 0.0279$). This result mimics the 1-hop codon-changes distribution found in Supp. Fig. 3A. This is expected because 1-hop codon-changes occur much more frequently than multi-hop codon-changes, thus enrichment at the amino-acid level is dominated by the effect seen at the 1-hop codon-level.
**(B)** Enrichment (log$_2$ fold-change) over model expectations for all 192 amino acid substitutions found in ALE. Mean value and standard deviation are shown for each distribution. ALE mutations show enrichment in synonymous mutations, but not to the extent found in WT variants.
**(C)** Enrichment (log$_2$ fold-change) over model expectations for all 160 amino acid substitutions found in LTEE. Mean value and standard deviation are shown for each distribution.

### Supp. Fig. 7: Pairwise comparisons of WT, ALE, and LTEE amino acid substitutions show relative differences in enrichment.

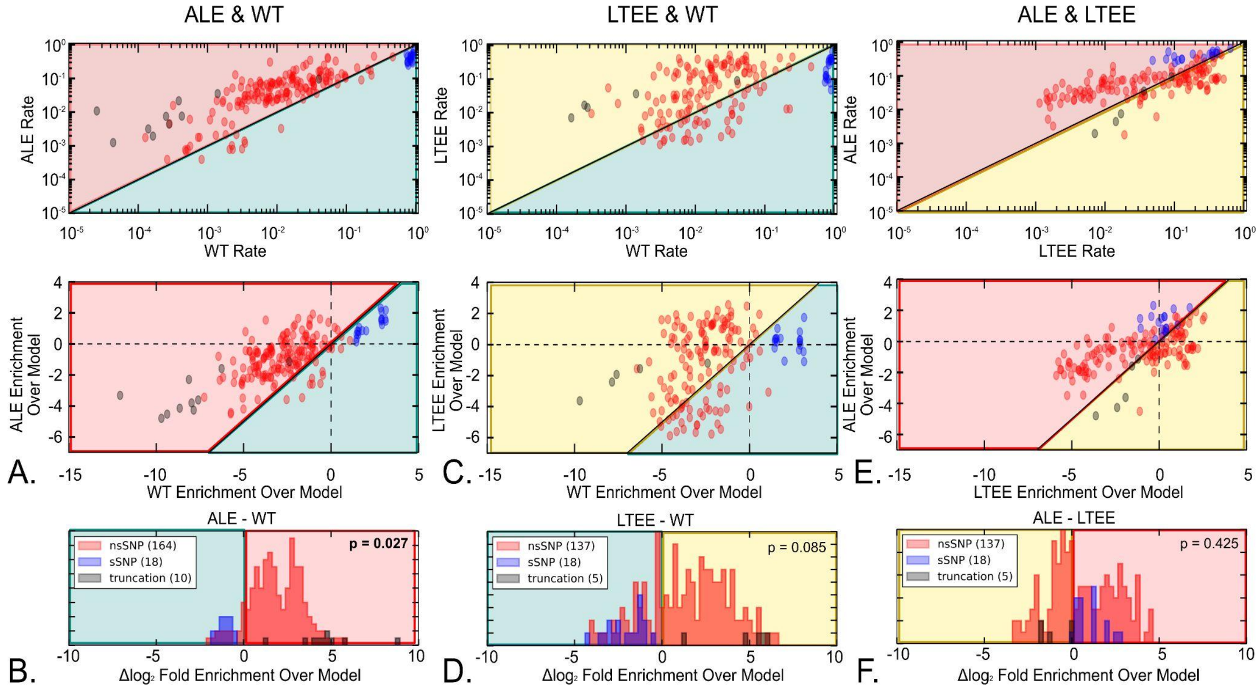


**(A)** Rate of occurrence 192 specific amino acid substitutions found in both ALE and WT data (top). The occurrence rates are normalized ($\log_2$-fold enrichment) using the expected value (bottom). Synonymous (blue) and non-synonymous (red) mutations are shown. Truncations are shown (gray).

**(B)** For each amino acid substitution, the relative change in enrichment is calculated. The distribution of this relative change is shown. Compared to ALE, WT mutations are more likely to fix synonymous mutations (p = 0.027).

**(C)** Rate of occurrence 160 specific amino acid substitutions found in both LTEE and WT data (top). The occurrence rates are normalized ($\log_2$-fold enrichment) using the expected value (bottom). Synonymous (blue) and non-synonymous (red) mutations are shown.

**(D)** For each amino acid substitution, the relative change in enrichment is calculated. The distribution of this relative change is shown. WT mutations are more likely to fix synonymous mutations relative to LTEE.

**(E)** Rate of occurrence 160 specific amino acid substitutions found in both ALE and LTEE data (top). The occurrence rates are normalized ($\log_2$-fold enrichment) using the expected value (bottom). Synonymous (blue) and non-synonymous (red) mutations are shown.

**(F)** For each amino acid substitution, the relative change in enrichment is calculated. The distribution of this relative change is shown.
